# Sexual segregation in a highly pagophilic and sexually dimorphic marine predator

**DOI:** 10.1101/472431

**Authors:** Christophe Barbraud, Karine Delord, Akiko Kato, Paco Bustamante, Yves Cherel

## Abstract

Sexual segregation is common in many species and has been attributed to intra-specific competition, sex-specific differences in foraging efficiency or in activity budgets and habitat choice. However, very few studies have simultaneously quantified sex-specific foraging strategies, at sea distribution, habitat use, and trophic ecology. Moreover, these studies come from low latitude areas reflecting a lack of evidence for polar species. We investigated sexual segregation in snow petrels *Pagodroma nivea* and combined movement, foraging trip efficiency, stable isotope and oceanographic data to test whether sexual segregation results from sex-specific habitat use. Breeding birds foraging in the Dumont d’Urville sea, Antarctica, were tracked during incubation. Some similarities between males and females foraging characteristics did not support the sexual segregation hypothesis. Indeed, space-use sharing and utilization distribution, δ^13^C values and foraging trip performances (trip duration, length, speed and directions, mass gain, proportion mass gain) were similar between males and females.. However, there was support for sexual segregation in foraging characteristics linked to foraging habitats. Females foraged less than males in areas with higher sea ice concentration (SIC >70%) and had lower δ^15^N values in plasma, blood cells and feathers. Foraging efficiency (proportionate daily mass gain while foraging), was greater for females than for males, and was greater for larger females with deeper bills. Females were more efficient than males during short (<2 days) foraging trips, and for females, but not for males, mass gain, proportion mass gain and body condition at return from a foraging trip were positively correlated to SIC of the foraging areas. Together, these results suggest an absence of sexual segregation at large spatial scales in snow petrels during incubation, but strongly support habitat segregation between high (>70%) more profitable SIC (males) and low SIC areas (females), probably driven by intraspecific competition. Therefore, male and female snow petrels segregate at small spatial scales mainly determined by habitat (SIC) characteristics.

## Introduction

Sexual segregation occurs in a many living animals including invertebrates (Hochkirch et al. 2007, Romey & Wallace 2007) and vertebrates (Ruckstuhl & Neuhaus 2005, Wearmouth & Sims 2008) but also in plants (Harder et al. 2000). Investigating sexual segregation is of particular relevance from a fundamental point of view to understand how and why the sexes differentially distribute themselves and the consequences on population processes and dynamics. It is also relevant from a management and conservation point of view since sex specific distribution influences overlap with spatial distribution of human activities and/or contamination gradient (Carravieri et al. 2014). Two main concepts have been proposed to describe sexual segregation: social segregation, where males and females tend to form single-sex groups within the same or homogeneous habitat; and habitat segregation, where males and females use different habitats within a home range and with habitats differing in their amount or quality of forage distributed heterogeneously or patchily (Conradt 2005, Ruckstuhl 2007). Both social segregation and habitat segregation can or cannot lead to spatial or temporal segregation, which have been be considered as auxiliary concepts (Conradt 2005, Ruckstuhl 2007). However, the distinction between habitat segregation and spatial segregation is scale-dependent: spatial segregation is a mechanism to avoid competing for the same habitat by choosing different locations, while habitat segregation is a mechanism to avoid competing for resources at the same location. These concepts can also be understood in the framework of equalizing and stabilizing mechanisms applied to movement ecology (Chesson 2000, Jeltsch et al. 2013).

Several hypotheses have been proposed to explain social and habitat segregations (Conradt 2005, Ruckstuhl 2007, Wearmouth & Sims 2008). In solitary animals, social segregation is unlikely to occur since by definition a single animal is not social (Conradt 1998, Neuhaus & Ruckstuhl 2004), except perhaps in rare cases (Martin & Da Silva 2004). Four main hypotheses explain habitat segregation in solitary species (Ruckstuhl 2007, Wearmouth & Sims 2008). The forage-selection hypothesis, which incorporates the scramble competition hypothesis, suggests sex differences in nutritional requirements linked to sex-specific differences in body size (Gross 1998). The larger sex individuals select habitats where intake rates are high whereas the smaller sex individuals are constrained to sites where they can obtain a high-quality food (Beier 1987, Barboza & Bowyer 2000). Alternatively, one sex may forage more efficiently, thus outcompeting and excluding the other (scramble-competition hypothesis or intersexual competition hypothesis) (Clutton-Brock et al. 1987). The activity-budget hypothesis, initially developed for group-living species, was extended to solitary species and to species with unequal reproductive investment (Wearmouth & Sims 2008). This hypothesis proposes that sex differences in activity budgets will increase with divergence in the body size of the sexes. Therefore, the sex-specific energy requirements will result in sex-specific habitat used due to allometric relationships between body size and metabolic rate. Finally, the predation-risk hypothesis proposes sexual differences in risk of predation and in reproductive strategies (Main et al. 1996), and the thermal niche-fecundity hypothesis assumes that sex differences occur in the temperature at which fecundity is maximized (Sims 2005).

Sexual segregation has been widely studied among terrestrial animals, particularly mammals, but only relatively recently in marine organisms (Wearmouth & Sims 2008). Yet, despite an ongoing interest in sexual segregation in marine animals such as seabirds and marine mammals (Lewis et al. 2002, Elliott et al. 2010, Phillips et al. 2011, Mancini et al. 2013, Baylis et al. 2016, Kernaléguen et al. 2016), the underlying causes and the mechanisms driving habitat segregation remain poorly understood. In addition, very few studies focused on between-sex differences in habitat segregation in relation to dynamic oceanographic features (Pinet et al. 2012, Cleasby et al. 2015, Paiva et al. 2017), thereby limiting our ability to distinguish between the concurrent sexual segregation hypotheses. Moreover, these studies come from temperate or tropical areas reflecting a lack of evidence for polar species. However, foraging strategies may differ between polar, temperate and tropical oceanographic environments, at least in seabirds (Baduini & Hyrenbach 2003, Weimerskirch 2007). Furthermore, for practical, technical and ethical reasons most studies that have investigated sexual segregation on marine animals have focused on large species (Phillips et al. 2011), complicating the possibility to discriminate between the various hypotheses proposed to explain sexual segregation.

In this study we aimed to quantify sexual differences in the foraging strategies, at sea distribution, habitat use, and trophic ecology of a sexually dimorphic polar seabird, the snow petrel, *Pagodroma nivea*, during the incubation period. Snow petrels are endothermic animals, therefore excluding the thermal niche-fecundity hypothesis as an explanatory hypothesis. Since predation on this species is occasional and no sex-specific predation is known to occur (Barbraud 1999), the predation-risk hypothesis can be discounted. Therefore, both the forage-selection hypothesis and the activity budget hypothesis can be highlighted as possible mechanisms for segregation in this species. There is considerable overlap between the forage-selection hypothesis and the activity-budget hypothesis, complicating our ability to make clear predictions to distinguish between the two, and to estimate the relative support of each hypothesis (Wearmouth & Sims 2008). Nevertheless, using GPS tracking data, isotopic data and environmental data we addressed the following main questions: (1) do female snow petrels differ from males in their foraging tactics, distribution and habitat use?; (2) how are body reserves regulated during incubation in the two sexes?; and (3) do sex-specific morphological characteristics influence foraging efficiency? Based on results from comparative studies suggesting that dimorphic seabird species from polar/temperate regions are more prone to show trophic or spatial segregation than dimorphic species from the tropics (Mancini et al. 2013), and on a relationship between sexual segregation in diet and sexual size dimorphism in seabirds (Phillips et al. 2011), we predicted sexual segregation in diet and/or spatial segregation in the snow petrel, which is one of the most sexually dimorphic seabird species (Croxall 1982, Fairbairn & Shine 1993).

## Methods

### Study species

The snow petrel is endemic to Antarctica and the Southern Ocean, with a circumpolar breeding distribution (Croxall et al. 1995). It is a specialist forager and ship-based observations indicate that this is the most pagophilic species amongst flying seabirds, occurring only where there is some degree of sea ice cover (Griffiths 1983, Ainley et al. 1984, 1986), generally within the marginal ice zone and areas of heavy ice concentrations (Ainley et al. 1992, 1993). Snow petrels forage by flying rapidly along the edges of ice floes, ice shelves and icebergs in search of its prey (Ainley et al. 1984). The species feeds primarily on fish, including the myctophid *Electrona antarctica* in oceanic waters and the pelagic nototheniid *Pleuragramma antarctica* (Antarctic silverfish) in neritic waters; they prey also upon swarming crustaceans, the Antarctic (*Euphausia superba*) and ice (*E. crystallorophias*) krill, and the hyperiid amphipod *Themisto gaudichaudii* (Ainley et al. 1984, 1991, Ridoux & Offredo 1989, Van Franeker & Williams 1992, Ferretti et al. 2001). At Pointe Géologie (Adélie Land), undetermined fish dominated the chick diet in 1982 (Ridoux & Offredo 1989) and fish items identified in 1994 were all Antarctic silverfish (authors’unpublished data). Prey are caught by dipping and surface-seizing (Harper et al. 1985) generally on the wing but also by ambush feeding (Ainley et al. 1984). Snow petrels breed in crevices and under boulders. Adult birds arrive at the colonies in late October to copulate before departing at sea for a two to three week pre-laying exodus, and females lay a single egg in early December (Mougin 1968, Isenmann 1970). Incubation lasts ≈44 days on average during which males and females alternately incubate their egg until hatching (Brown 1966, Barbraud et al. 1999). After hatching the chick is guarded by parents alternating short spells until it attains homeothermy. Then the chick is left unattended and regularly fed by both parents until fledging, which occurs on average ≈47 days after hatching. Adults leave the colony during the first two weeks of March before dispersing at sea where they remain in the sea ice zone during the non-breeding period (Delord et al. 2016).

### Fieldwork

Fieldwork was carried out at Ile des Pétrels (66°40’S, 140°01’E), Pointe Géologie archipelago, Adélie Land, East Antarctica, between 7 December 2015 and 17 January 2016. This corresponds to the incubation period. On average 550 pairs of snow petrels breed on Ile des Pétrels in dense colonies or in loosely aggregated nests (CEBC-CNRS unpublished data). By daily visits at 36 nests, we studied laying dates and the duration of the foraging trips and incubation shifts of 36 males and 36 females until hatching. Incubating birds were identified using their metal ring number. Sixty five snow petrels (n = 36 females and n = 29 males) were tracked with GPS loggers (nanoFix-Geo; PathTrack Limited, UK) during the incubation period. We tracked only one foraging trip per bird to minimize disturbance and to ensure independence between trips. The devices weighed 2.2 g, which represented between 0.5% and 0.8% of the birds’ mass, thus well below the 3% threshold advised by Phillips et al.(2003). Birds were manually captured at the nest and weighted (± 5 g) in a bag with a Pesola spring balance before being equipped with a GPS. The birds were initially sexed by vocalization when approached on the nest and handled (male calls have a lower pitch and a lower rhythm than those of females (Guillotin & Jouventin 1980, Barbraud et al. 2000). GPS units were deployed on birds about to leave for a foraging trip (i.e. when both partners were at the nest) and were attached to the two central tail feathers using Tesa^®^ tape. The GPS recorded locations at 15, 30, 40 or 60 min intervals. Several intervals (15 min, n = 15; 30 min, n = 4; 40 min, n = 43; 60 min, n = 3) were tested to estimate the minimum interval frequency that allowed the GPS battery to last for a complete foraging trip. Birds were recaptured on the day they returned to the nest following their foraging trip, weighed, measured (wing length ± 1 mm with a ruler, tarsus length, bill length, and bill depth ± 0.1 mm with calipers) and the loggers were recovered. All birds were recaptured but three birds lost their GPS during the foraging trip. Data from all other GPS (n = 62) were retrieved successfully.

### Tissue sampling, molecular sexing and stable isotopes

Adults equipped with GPS and 24 additional individuals (11 females and 13 males) were sampled during incubation for stable isotope and molecular sexing analyses. A blood sample from the alar vein was taken immediately after capture of the bird upon return from a foraging trip using a 1-mL heparinized syringe and a 25-gauge needle and maintained at 4°C until being processed. Collected blood volumes ranged from 0.50 to 0.80 mL. Blood samples were separated into plasma and blood cells by centrifugation at 12,000 rpm for 5 min, within 2-3 hours of sampling and stored frozen at −20°C until analyses at the laboratory. For each individual, 6 whole body feathers were pulled out from the upper chest and stored dry in sealed individual plastic bags for stable isotope analysis.

From a subsample of blood cells, the sex was determined by polymerase chain reaction amplification of part of two highly conserved genes present on the sex chromosomes as detailed in Weimerskirch et al. (2005). Stable carbon (δ^13^C) and nitrogen (δ^15^N) isotope ratios in the blood cells, plasma and body feathers of snow petrels were determined to investigate the trophic choices of each sex and consistency of their foraging niche over time. The isotopic method was validated in the Southern Ocean for several seabird species: δ^15^N values mainly define the trophic position, with values increasing with trophic level (Cherel et al. 2010), and δ^13^C values indicate the latitude of the foraging habitat (Cherel & Hobson 2007, Jaeger et al. 2010). Plasma has a half-life of about 3 days (Hobson & Clark 1993), a shorter period than the average trip duration during incubation (≈7 days, Barbraud et al. 1999), and represents prey ingestion and trophic ecology during the last trip before sampling (Cherel et al. 2005a). Blood cells have a half-life of about 30 days (Hobson & Clark 1993) and represent dietary information integrated over a few months. Feathers contain dietary information at the time they were grown, because keratin is inert after synthesis (Hobson & Clark 1992, 1993, Bearhop et al. 2002). In snow petrels body moult is a gradual process extending over at least 4 months in summer and autumn. It begins during incubation, but most body feathers grow in the weeks following completion of breeding, i.e. from February to April (Maher 1962, Beck 1970). Therefore, isotopic values of body feathers contain information about diet near the end of the previous breeding season and the beginning of the previous non-breeding season.

Feathers (one single feather per bird) were cleaned to remove surface contaminants using a 2:1 chloroform:methanol solution followed by two methanol rinses. They were then oven dried for 48 h at 50°C and cut into small pieces using stainless steel scissors. Blood cells and plasma samples were freeze-dried and powdered. Since avian plasma, unlike blood cells, contains a high and variable lipid content that affect its δ^13^C values, lipids were removed from plasma samples using chloroform/methanol (Cherel et al. 2005a, Cherel et al. 2005b). Then, tissue sub-samples were weighed with a microbalance (aliquots mass: ≈ 0.3 mg dw), packed in tin containers, and nitrogen and carbon isotope ratios were subsequently determined at the laboratory LIENSs by a continuous flow mass spectrometer (Thermo Scientific Delta V Advantage) coupled to an elemental analyser (Thermo Scientific Flash EA 1112). Results are presented in the usual δ notation relative to Vienna PeeDee Belemnite and atmospheric N_2_ for δ^13^C and δ^15^N, respectively. Replicate measurements of internal laboratory standards (acetanilide and peptone) indicate measurement errors <0.15 ‰ for both δ^13^C and δ^15^N values.

### Foraging analysis and spatial usage

Spatial and statistical analyses were performed using R 3.2.1 using the “stats” package (R Development Core Team 2015) and “*adehabitatLT*” package (Calenge 2006, Calenge et al. 2009). From the GPS recorded data, foraging trips were reconstructed and data were rediscretized to have one location each 40 min. Some of the trips were largely incomplete (return journey not initiated; n = 15 corresponding to the 15 min intervals) because of battery limitations and were removed from the analysis. For each complete (n = 40) and incomplete (return journey initiated; n = 7) foraging trip, we computed the following foraging indices: maximum distance to the colony (*Dmax*, km), average movement speed (*MS*, km h^−1^) and daily distance covered (*Dday*, km d^−1^). For each complete trip, we calculated the additional following metrics: total distance travelled (*Dtotal*, km) and trip duration (*T*, h). Spatial distribution of snow petrels was investigated by producing utilization distributions (UDs 25%, 50%, 75% and 95%; Worton 1989) for each individual, using kernel analysis with a cell size of 0.1° × 0.1° and a smoothing parameter (h) that was estimated using the ad hoc method href. Grid cell size was based on the mean accuracy of the devices (≈ 10 m), the mean maximum speed of flying snow petrels (see Results) and on the time interval between two GPS locations (40 min). To investigate whether space use differed between sexes, we calculated observed overlaps in each UD representing the high core (25%), core (50%), middle (75%) and general (90%) use areas using utilization distribution overlap index (UDOI), which is the most appropriate measure of quantifying similarity among UD estimates (Fieberg & Kochanny 2005). The extent of overlap between male and female home ranges was estimated using Bhattacharyya’s affinity (BA), which ranges from 0 (no overlap) to 1 (complete overlap). Using these metrics we performed a randomization procedure to test the null hypothesis that there was no difference in the spatial distribution of males and females at the population level (Breed et al. 2006). The sex of each bird was randomly assigned using the observed sex ratio in our data set and the overlap metric between males and females was calculated for 25%, 50%, 75% and 95% kernels. We performed 1000 randomizations of our dataset from which the probability of accepting the null hypothesis was calculated as the proportion of random overlaps that were smaller than the observed overlap. Since we were testing only if the observed overlap was smaller than random overlap, we considered this as a one-tailed test. Second, we tested the null hypothesis that there was no difference in the extent of overlap in spatial distribution of males and females at the individual level.

For each foraging trip we also calculated the following metrics from the phenotypic data: the body mass change (Δm, in g) between departure and arrival of a foraging trip, the daily mass gain (Mday, in g.day^−1^) calculated as the ratio between Δm and the trip duration, the proportion mass gain calculated as the ratio between Δm and mass at departure for a foraging trip, and the proportion daily mass gain calculated as the ratio between Daym and mass at departure for a foraging trip. A body condition index before departure and after return from a foraging trip was also calculated. To estimate the body condition we used the body measurements to calculate the scale mass index (SMI) as recommended by Peig and Green (2009, 2010). The SMI adjusts the mass of all individuals to that expected if they had the same body size. We used the score of the first axis of a principal component analysis (PC1) combining wing, bill, tarsus lengths and bill depth to characterize body size. PC1 accounts for 70.9% of the total variance and all measurements are highly correlated with PC1 (Pearson’s r > 0.80; P < 0.001). The SMI was calculated for each individual *i* according to the formula:

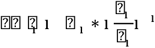

where *M_i_* and *L_i_* are, respectively, the body mass and the PC1 score of the individual *i, L_0_*, is the value of PC1 for the whole studied population and *b* the slope estimate of the RMA (Reduced Major Axis) regression of log-transformed body mass on log-transformed PC1.

### Foraging habitat covariates

To investigate the foraging habitats used by males and females, the tracking locations were categorized as occurring during commuting (outward and inward) or foraging (middle) stage of foraging trips, as commonly used for central-place foragers. Among Procellariforms the distinction between these stages varies greatly between species and breeding stages (Weimerskirch et al. 1997, Phillips et al. 2009). Moreover, at the individual level defining objectively the transition between such behaviors may prove to be difficult (Phillips et al. 2009, Wakefield et al. 2009). To avoid this pitfall, we applied the method used by Wakefield et al. (2009) and Phillips et al. (2009) to determine the stage of the trips at which the transitions occurred at the population level. For each location within a foraging trip the ratio *d_col_/D_max_* was calculated, where *d_col_* is the distance from the colony and *D_max_* is the maximum distance from the colony reached during that trip. The ratio *t/T* was also calculated, where *t* is the time elapsed since the beginning of the trip and *T* is the total trip time. Then, the total variance in *d_col_/D_max_* for all locations occurring before *t/T* was plotted against *t/T*. The point of inflexion of this curve was determined as well as the value of *t/T* at this point. Tracking locations recorded before this point were classified as those corresponding to the outward trip. Similarly, the total variance in *d_col_/D_max_* occurring after *t/T* was plotted against *t/T* and the *t/T* value from which a monotonic decrease of the variance began was recorded. Tracking locations recorded after this point were classified as those corresponding to the return trip, and locations between both points were considered as foraging locations.

Previous studies have shown that the snow petrel is a sea ice obligate species and remains highly associated with sea ice year round (Griffiths 1983, Ainley et al. 1984, 1986, 1992, 1993, Delord et al. 2016). We therefore used sea ice concentration (SIC) to describe the foraging habitat of snow petrels. Although sea surface temperature is commonly used to describe foraging habitats in seabirds, there are very few sea surface temperature observations in regions covered by sea ice, especially in the Southern Ocean (Rayner et al. 2003). Therefore this covariate could not be used. We used passive-microwave estimates of daily sea ice concentration from the Special Sensor Microwave Imager (SSMI/I) brightness temperatures (12.5 × 12.5 km resolution) from the Institut Français de Recherche pour l’Exploitation de la Mer (Ifremer, ftp://ftp.ifremer.fr/ifremer/cersat/products/gridded/psi-concentration/data/antarctic). We also used bathymetry data (ocean depth at one-minute horizontal spatial resolution) obtained from NOAA’s ETOPO (https://sos.noaa.gov/datasets/etopo1-topography-and-bathymetry/) as an additional habitat variable. Daily sea ice concentration and depth values were extracted for each foraging location (therefore excluding the commuting part of the trips at sea) on each track using bilinear interpolation from the native ice and depth grids using “*raster*” package in R (Hijmans 2018). Since snow petrels are highly associated with the sea ice region (as defined by the region within >15% sea ice concentration isocline, Cavalieri et al. 1991), the SIC data were filtered to retain SIC values >15%.

### Statistical analysis

Isotopic niche of the two sexes was established using the metric SIBER (Stable Isotope Bayesian Ellipses), which is based on a Bayesian framework that confers a robust comparison to be made among data sets concerning different sample sizes (Jackson et al. 2011). The area of the standard ellipse (SEA_C_, an ellipse having a 40% probability of containing a subsequently sampled datum) was used to compare female and male isotopic values and their overlap in relation to the total niche width (i.e. both sexes combined), and a Bayesian estimate of the standard ellipse and its area (SEA_B_) was used to test whether females’ isotopic niche is narrower than males’ isotopic niche (Jackson et al. 2011). The *standard.ellipse* and *convexhull* functions were used to calculate these metrics from SIBER implemented in the package ‘*SIAR*’ (Parnell et al. 2010) under R. Consistency in foraging niche was estimated following Votier et al. (2010) and Ceia et al. (2012), by regressing stable isotope ratios in plasma on those of blood cells to obtain an index of consistency in carbon source (habitat) and trophic level. Since δ^13^C has a trophic component, we used the studentized residuals of the relationship with δ^15^N in the same tissue (male plasma: F_1,25_=1.438, P=0.242, r=0.233; male blood cells:: F_1,36_=0.838, P=0.366, r=0.151; female plasma: F_1,33_=1.470, P=0.234, r=0.206; female blood cells: F_1,45_=6.507, P=0.014, r=0.355) to determine the degree of short-term repeatability in δ^13^C independently of trophic effects. Longer-term foraging consistency was estimated by regressing stable isotope values of blood cells (actual breeding period) with those of feathers (most likely the end of the previous breeding period and subsequent fall at sea). We also used the residuals to correct the trophic component associate with δ^13^C by regressing these values upon δ^15^N signatures in feathers (male: F_1,36_=1.945, P=0.172, r=0.226; female: F_1,45_=5.863, P=0.020, r=0.340).

Foraging probability was modelled using a binomial generalized additive mixed model (GAMM) in the ‘*gamm4*’ package in R (Wood et al. 2017). This allowed for the possibility of nonlinear responses to environmental covariates, which we expected. The response variable was the tracking location, which was coded as 1 for a foraging location and as 0 for a commuting location, and explanatory variables were sea ice concentration and bathymetry. Because interactions between the variable sex and environmental covariates would be difficult to interpret in complex nonlinear models, separate models were developed for male and female birds. Models included sea ice concentration and bathymetry as fixed factors, and bird identity as a random term to account for pseudoreplication issues. The smoothing parameter was chosen automatically using generalized cross-validation. To model spatial auto-correlation an isotropic thin plate spline was included, set up as a two dimensional smoother based on both x and y coordinates (Cleasby et al. 2015). To ascertain whether collinearity between covariates may have occurred we examined the correlations between environmental variables using a Spearman correlation coefficient since covariates were not normally distributed. We assumed that a correlation of greater than r_s_ 0.4 was problematic, but the correlation was below this threshold (r_s_ = 0.14).

Foraging intensity was modelled using GAM in the ‘*gam*’ package in R (Hastie & Tibshirani 1990). Foraging intensity was defined based on the frequency distribution of the tracking locations classified as foraging only. The environmental covariates were divided into K classes. Then, within each class the number of foraging locations was extracted and the count was used as the response variable. A GAM with a quasi-Poisson distribution was then fitted to the data. Separate models were developed for male and female birds. Models included sea ice concentration and bathymetry as fixed factors. The smoothing parameter was chosen automatically using generalized cross-validation. For SIC we used K=1% SIC classes and for bathymetry we used K=50 m classes.

Differences between sexes in body measurements, sea ice characteristics used, and foraging trip metrics were tested using Student’s t-tests and differences in stable isotope date using Student’s t-tests and Wilcoxon rank tests. Since we performed a large number of tests when comparing male and female body size measurements, isotopic values and foraging trip characteristics, we used the Benjamini-Hochberg procedure (Benjamini & Hochberg 1995) to control for false discovery rate. We chose a false discovery rate (q*) of 0.10 when applying the Benjamini-Hochberg procedure. This choice was motivated by the fact that this study was conducted on a single year and was exploratory. In such cases setting a FDR to an extremely low value results in decreasing the statistical power for detecting genuine effects and several authors recommend setting FDR to a relatively large value (Yoccoz 1991, Field et al. 2004, Roback and Askins 2005).

## Results

Male snow petrels were larger than females, particularly for bill length and bill depth, and were 10% heavier than females (Table 1). Bill length, bill depth and body mass were the most sexually dimorphic phenotypic traits.

**Table 1.** Body measurements of male and female snow petrels and percentage of difference between sexes for each measurement. For t-tests homogeneity of variances we checked using a Brown and Forsythe tests (Brown and Forsythe 1974), and corrected p values are reported (uncorrected in brackets). Significant differences with a false detection rate of 0.10 are shown in bold. Δ is the difference in %. The sample size is 47 individuals.

| | Sex | Mean (SD) | Range | $\Delta$ | t-test |
| --- | --- | --- | --- | --- | --- |
| Wing length (mm) | Male | 298.3 (6.8) | 287.0-311.0 | 2.3 | **$t_{45}=4.215$** |
|  | Female | 290.0 (6.7) | 280.0-302.0 |  | **$p<0.001$** (<0.001) |
| Tarsus length (mm) | Male | 40.2 (1.2) | 38.0-42.7 | 4.5 | **$t_{45}=4.846$** |
|  | Female | 38.4 (1.2) | 35.9-41.0 |  | **$p<0.001$** (<0.001) |
| Bill length (mm) | Male | 24.4 (1.0) | 22.1-26.6 | 9.0 | **$t_{45}=8.488$** |
|  | Female | 22.2 (0.8) | 20.9-23.7 |  | **$p<0.001$** (<0.001) |
| Bill depth (mm) | Male | 10.8 (0.5) | 9.6-11.8 | 8.3 | **$t_{45}=6.408$** |
|  | Female | 9.9 (0.4) | 8.9-10.9 |  | **$p<0.001$** (<0.001) |
| Body mass<sup>1</sup> (g) | Male | 389.0 (30.4) | 331.5-464.0 | 10.3 | **$t_{45}=6.014$** |
|  | Female | 340.5 (24.0) | 300.0-385.0 |  | **$p<0.001$** (<0.001) |
| Body mass<sup>2</sup> (g) | Male | 431.4 (34.0) | 365.0-495.0 | 10.3 | **$t_{45}=4.467$** |
|  | Female | 387.0 (33.9) | 320.0-460.0 |  | **$p<0.001$** (<0.001) |

<sup>1</sup>before a foraging trip; <sup>2</sup>on return from a foraging trip.

### Spatial distribution of males and females and habitat differences

Males and females foraged in offshore waters to the east and to the west of the colony in equal proportions (χ^2^=0.03, p=0.86, Figure 1). Space-use sharing was similar between males and females as the UDOI was not significantly lower than the null expectation for 25%, 50%, 75% or 95% UDs (Table 2). The 95% UDOI was > 1, indicating a higher than normal overlap between male and female UDs relative to uniform space use, i.e. male and female UDs were non-uniformly distributed and had a high degree of overlap. By contrast, the 25% UDOI was relatively close to 0 indicating less overlap between male and female UDs relative to uniform space use. Males and females UDs were also similar whatever the UDs considered since BA were not significantly lower than the null expectation for 25%, 50%, 75% or 95% UDs (Table 2).

**Figure 1.**
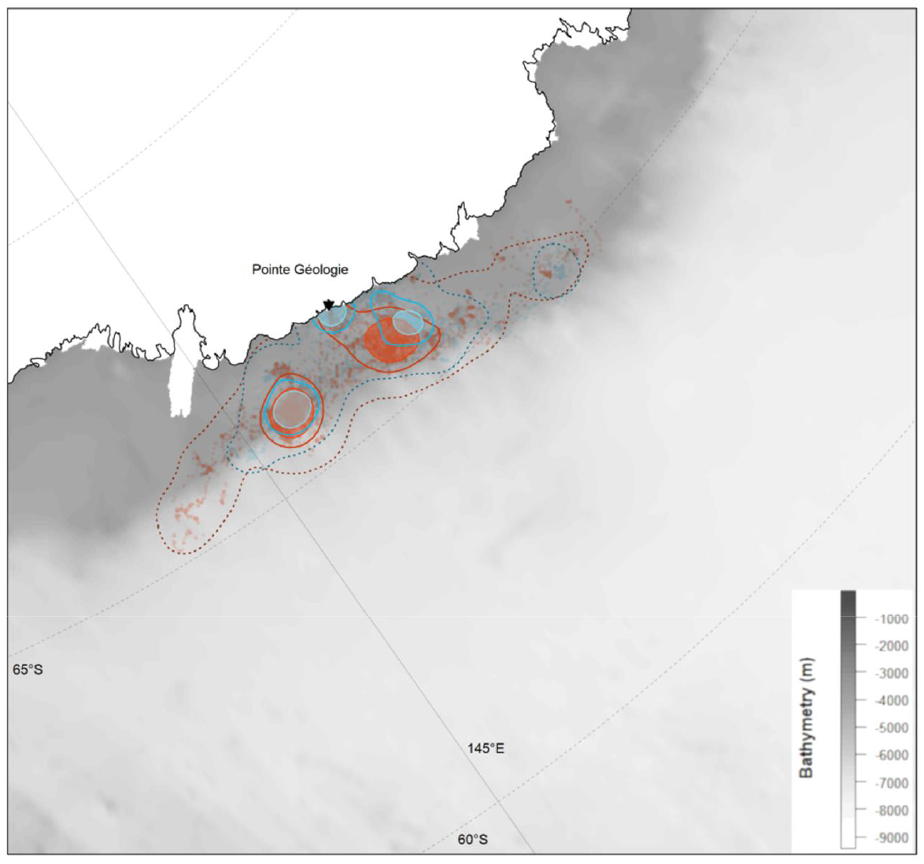
Foraging ranges of male (blue) and female (red) snow petrels during incubation sampled at Ile des Pétrels, Adélie Land, East Antarctica. Dots show raw location data. Kernel density based utilization distributions at 95% (dotted lines), 50% (solid lines) and 25% (filled areas). Bathymetry shown in grey and land in white. Ile des Pétrels is shown as a black triangle.

**Table 2.** Estimated overlap in utilization distributions (UD) between male and female snow petrels from Ile des Pétrels, Adélie Land, East Antarctica. UDOI: Utilization distribution overlap index. BA: Bhattacharyya’s affinity

| UD (%) | Observed UDOI | Randomized UDOI | p | Observed BA | Randomized BA | p |
| --- | --- | --- | --- | --- | --- | --- |
| 25 | 0.062 | 0.064 | 0.417 | 0.127 | 0.123 | 0.452 |
| 50 | 0.226 | 0.243 | 0.262 | 0.379 | 0.398 | 0.273 |
| 75 | 0.502 | 0.545 | 0.227 | 0.634 | 0.662 | 0.212 |
| 95 | 1.215 | 1.229 | 0.439 | 0.858 | 0.863 | 0.359 |

In average males foraged in areas with higher SIC than females (Table 3). Fitted models on foraging probability contained sex-specific smoothers for bathymetry and SIC (Table 4). For females, the GAMM model explained 10% of the deviance of foraging probability. All smoothers for SIC and bathymetry were significant (Table 4). Foraging probability increased sharply with increasing SIC up to 30% and more smoothly for high SIC (Figure 2). Foraging probability showed a first peak at depth of ≈600 m and a second and high peak at depth of ≈1600 m. Foraging probability sharply increased at depths >2500 m but sample size was small and there was high uncertainty. Both the random intercept for bird identity and the spatial smoother were significant.

**Figure 2.**
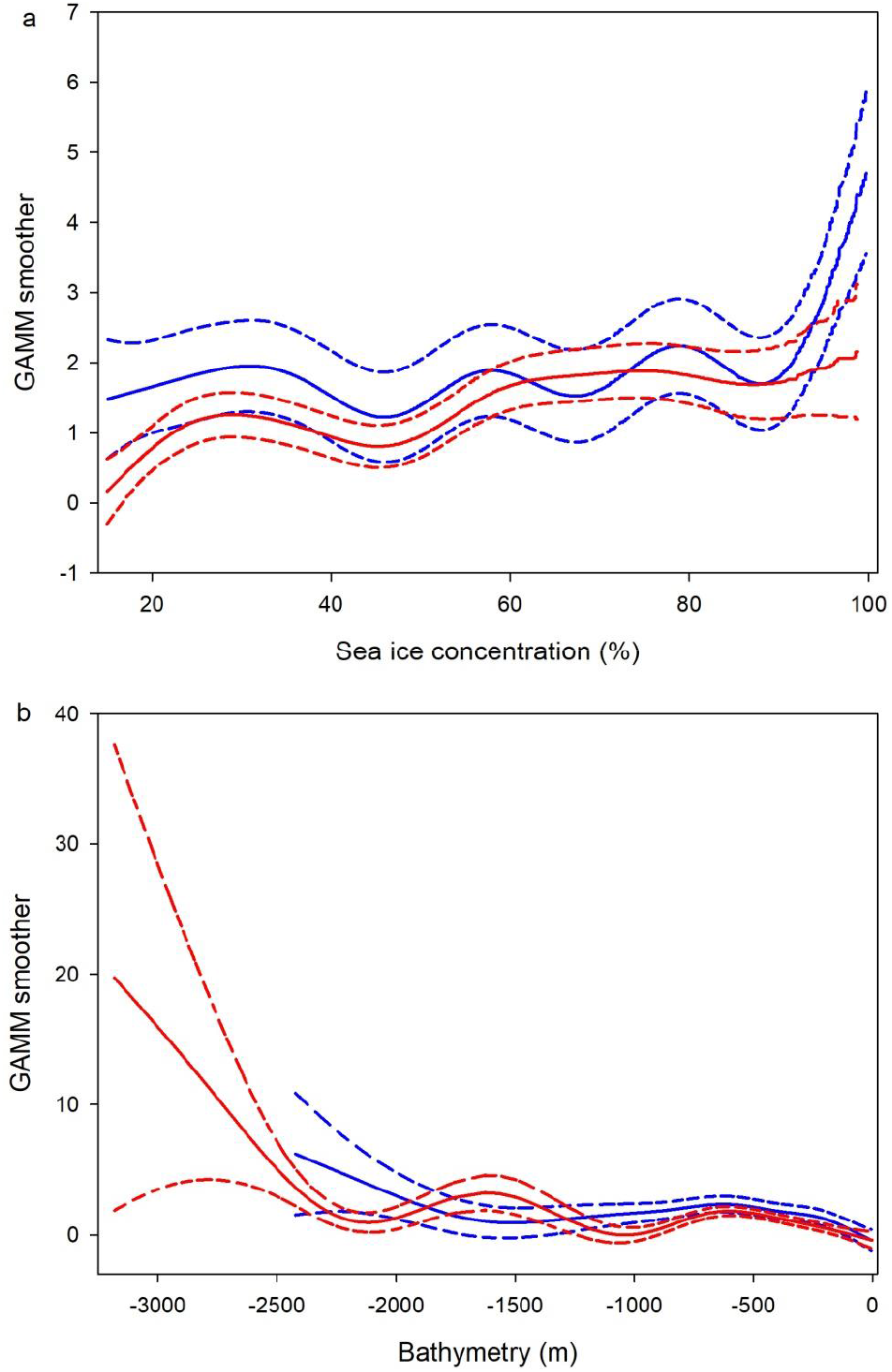
Foraging probability habitat selection functions for (a) sea ice concentration and (b) bathymetry). Plots show the predicted curve from the model (solid line) and 95% confidence intervals (dashed lines) for male (blue) and female (red) snow petrels sampled at Ile des Pétrels, Adélie Land, East Antarctica. GAMM: generalized additive mixed model.

**Table 3.** Mean, maximum and variance in sea ice concentration, mean, minimum and maximum bathymetry for foraging localities of male and female snow petrels from Ile des Pétrels, Adélie Land, East Antarctica. For t-tests homogeneity of variances we checked using a Brown and Forsythe tests (Brown and Forsythe 1974), corrected p values are reported (uncorrected in brackets). Significant differences with a false detection rate of 0.10 are shown in bold.

|  | Sex | Mean (SD) | Range | t-test |
| --- | --- | --- | --- | --- |
| Mean SIC (%) | Male | 54.8 (14.6) | 30.4-77.9 | **$t_{44}=2.600$** |
|  | Female | 44.0 (13.4) | 16.8-72.5 | **$p=0.063$** (0.013) |
| Maximum SIC (%) | Male | 83.8 (16.8) | 43.9-99.9 | $t_{44}=1.867$ |
| | Female | 72.6 (23.8) | 19.8-100.0 | $p=0.171$ (0.069) |
| Variance in SIC | Male | 375.7 (224.2) | 69.5-840.2 | $t_{44}=1.772$ |
| | Female | 264.0 (198.3) | 3.9-749.9 | $p=0.139$ (0.083) |
| Mean bathymetry (m) | Male | 503.8 (248.7) | 285-1490 | $t_{45}=0.911$ |
| | Female | 582.4 (341.2) | 259-1706 | $p=0.367$ (0.367) |
| Maximum bathymetry (m) | Male | 1354.1 (632.1) | 809-2973 | $t_{45}=1.285$ |
| | Female | 1617.4 (771.4) | 613-3223 | $p=0.257$ (0.205) |

**Table 4.** Generalized Additive Mixed Model (GAMM) results for foraging probability of male and female snow petrels as a function of sea ice concentration (SIC), bathymetry (BAT) and spatial autocorrelation (s(x,y)). edf indicates the estimated degrees of freedom.

| Variable | Sex | Smoother edf (p value) | Estimate (SE) | $\sigma^2$ (SE) |
| --- | --- | --- | --- | --- |
| Intercept | Male |  | 3.543 (1.101) |  |
|  | Female |  | 3.087 (1.365) |  |
| SIC | Male | **7.97 (<0.001)** |  |  |
|  | Female | **1.00 (0.003)** |  |  |
| BAT | Male | **3.89 (<0.001)** |  |  |
|  | Female | **1.00 (0.044)** |  |  |
| $s(x,y)$ | Male | **22.96 (<0.001)** | | |
|  | Female | **24.52 (<0.001)** |  |  |
| Random intercept for bird ID | Male |  |  | 10.250 (3.201) |
|  | Female |  |  | 1.174 (0.343) |

For males, the model explained 4.6% of the deviance of foraging probability. All smoothers for SIC and bathymetry, the random intercept for bird identity and the spatial smoother were significant (Table 4). Male foraging probability varied non-linearly with SIC and bathymetry. It increased smoothly with increasing SIC, and was higher when SIC was higher than ≈90% (Figure 2). Foraging probability also increased with bathymetry up to ≈600 m and remained relatively stable until ≈2000 m from which it increased.

Female foraging intensity was non-linearly related to SIC and bathymetry (Table 5). Foraging intensity increased with SIC up to a maximum for SIC ≈40% and then decreased for higher SIC (Figure 3). Lowest foraging intensity was observed for SIC >80%. Foraging intensity showed a rather bimodal distribution as a function of bathymetry. It was maximal in waters ≈400 m deep, then decreased to reach a minimum at ≈1400 m, and increased again for water depths between ≈2000-2700 m. Male foraging intensity was non-linearly related to SIC and bathymetry (Table 5). It showed a bimodal distribution as a function of SIC, with a maximum for SIC ≈36% and a second peak for SIC ≈85% (Figure 3). As for females, male foraging intensity was bimodal as a function of bathymetry. It was maximal in waters ≈400 m deep, then decreased to reach a minimum at ≈1500 m, and increased up to a second peak in waters ≈2400 m deep.

**Figure 3.**
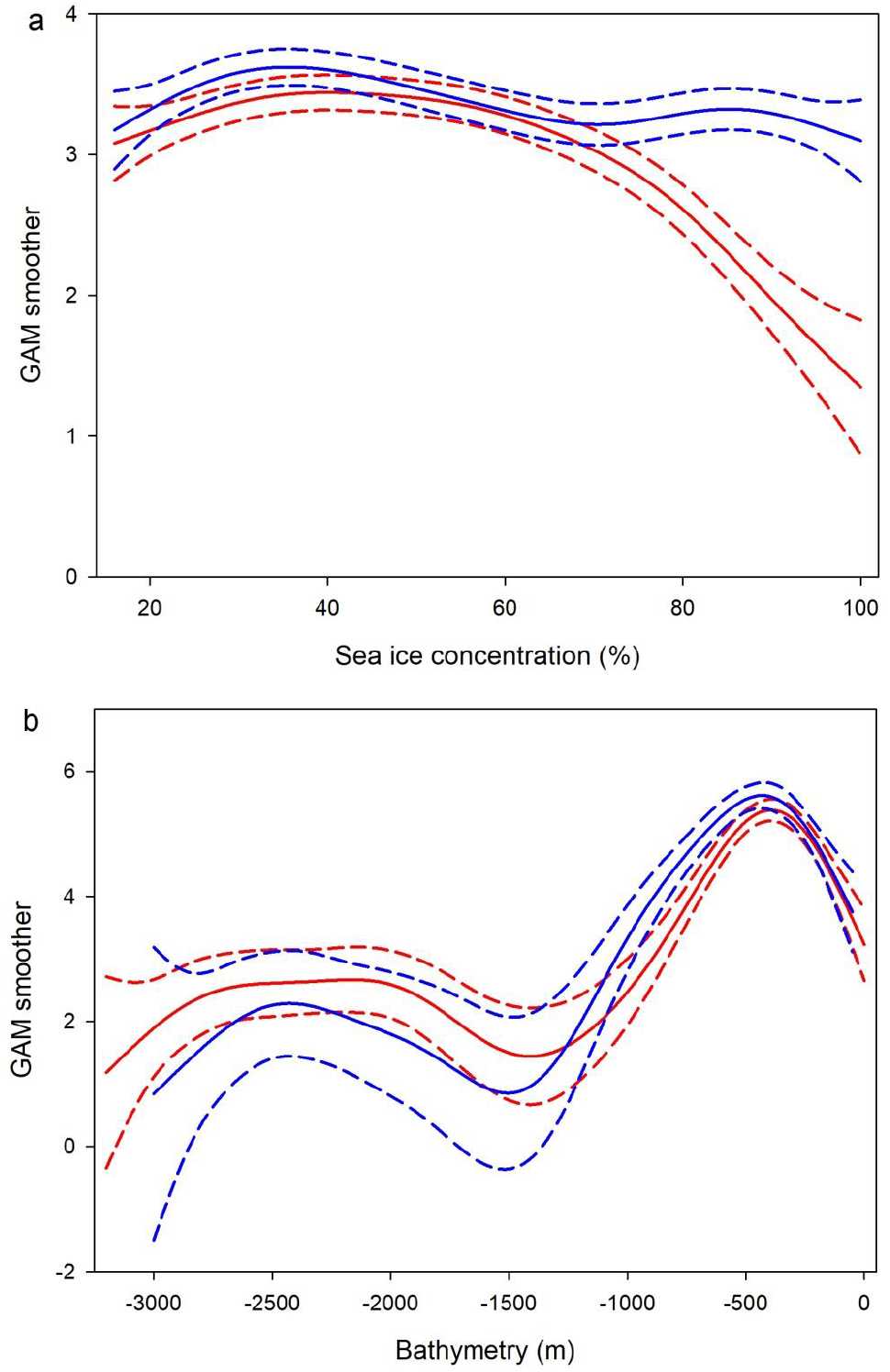
Foraging intensity habitat selection functions for (a) sea ice concentration and (b) bathymetry). Plots show the predicted curve from the model (solid line) and 95% confidence intervals (dashed lines) for male (blue) and female (red) snow petrels sampled at Ile des Pétrels, Adélie Land, East Antarctica. GAM: generalized additive model.

**Table 5.** Generalized Additive Model (GAM) results for foraging intensity of male and female snow petrels as a function of sea ice concentration (SIC) and bathymetry (BAT). edf indicates the estimated degrees of freedom.

| Variable | Sex | Smoother edf (p value) | Scale | Adjusted R<sup>2</sup> |
| --- | --- | --- | --- | --- |
| SIC | Male | **4.99 (<0.001)** | 2.50 | 0.223 |
|  | Female | **4.11 (<0.001)** | 2.25 | 0.663 |
| BAT | Male | **6.47 (<0.001)** | 16.01 | 0.779 |
|  | Female | **6.82 (<0.001)** | 9.28 | 0.863 |

### stable isotope ratios

Male plasma, blood cells and feathers had significantly 0.6-0.8‰ higher δ^15^N values than those of females (Table 6). There was no difference in δ^13^C values between males and females, except for plasma for which males had higher values. Males and females had similar SEA_B_ for all tissues (Figure 4). Overlap between SEA_B_ areas for males and females was 0.462, 0.586 and 0.599 for blood cells, plasma and feathers, respectively.

**Figure 4.**
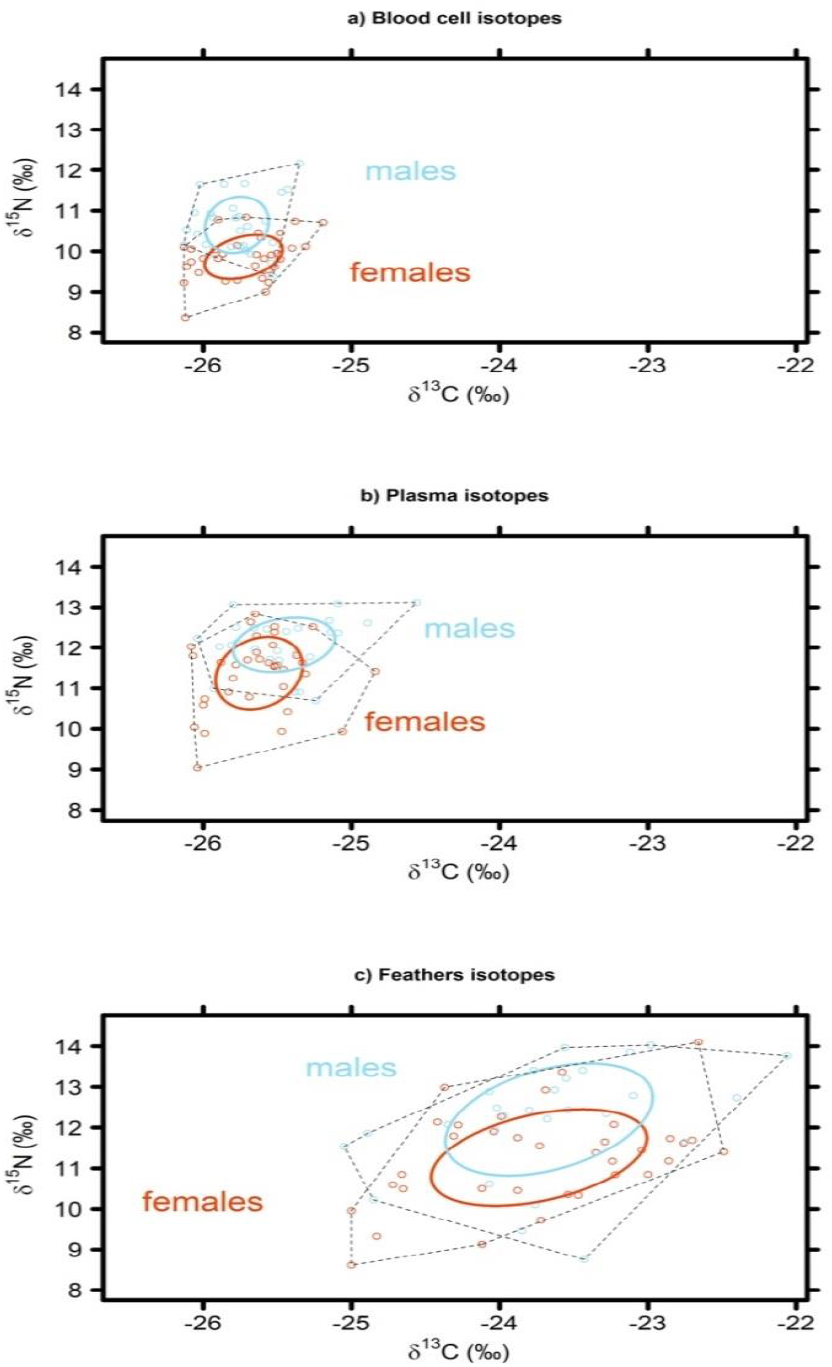
Isotopic niche area based on stable isotope values (δ^15^N and δ^13^C) in blood cells (top), plasma (middle) and body feathers (bottom) of male (blue) and female (red) snow petrels breeding at Ile des Pétrels, Pointe Géologie, Antarctica during the incubation period. The areas of the standard ellipses are represented by the solid lines, and the layman metric of convex hull area by black dotted lines.

**Table 6.** Stable isotope values in blood cells, plasma and feathers of male and female snow petrels sampled Ile des Pétrels, Adélie Land, East Antarctica. Values are mean ± SD. SEA_B_ are Bayesian approximation of the standard ellipse area. Values in brackets indicate n and range for δ^13^C and δ^15^N, and 95% credible interval for SEA_B_. p indicates the probability that SEA_B_ of males and females differ. Sample sizes are 47 individuals for blood cells and feathers, and 46 individuals for plasma. For t-tests homogeneity of variances we checked using a Brown and Forsythe tests (Brown and Forsythe 1974), corrected p values are reported (uncorrected in brackets). Significant differences with a false detection rate of 0.10 are shown in bold.

| Tissue | $\delta^{13}\text{C}(\text{‰})$ | | | $\delta^{15}\text{N}(\text{‰})$ | | | SEA<sub>B</sub> | | p |
| --- | --- | --- | --- | --- | --- | --- | --- | --- | --- |
|  | Male | Female |  | Male | Female |  | Male | Female |  |
| Plasma | -25.46 ± 0.35<br>(-26.01;-24.56) | -25.66 ± 0.22<br>(-26.06;-25.26) | **t<sub>83</sub>=2.105**<br>**p=0.057**<br>(0.039) | 12.11 ± 0.74<br>(10.70;13.38) | 11.49 ± 0.97<br>(9.04;12.83) | **t<sub>83</sub>=3.418**<br>**p=0.002**<br>(0.001) | 0.69<br>(0.45;0.99) | 0.79<br>(0.55;1.09) | 0.311 |
| Blood cells | -25.79 ± 0.21<br>(-26.13;-25.35) | -25.72 ± 0.27<br>(-26.13;-25.19) | t<sub>83</sub>=1.491<br>p=0.168<br>(0.140) | 10.65 ± 0.68<br>(9.37;12.17) | 9.96 ± 0.65<br>(8.36;11.46) | **t<sub>83</sub>=5.568**<br>**p<0.001**<br>(<0.001) | 0.41<br>(0.29;0.55) | 0.39<br>(0.29;0.52) | 0.586 |
| Feather | -23.68 ± 0.71<br>(-25.05;-22.06) | -23.50 ± 0.67<br>(-25.00;-22.49) | t<sub>83</sub>=0.335<br>p=0.738<br>(0.738) | 12.11 ± 1.35<br>(8.75;14.03) | 11.34 ± 1.35<br>(8.61;14.10) | **t<sub>83</sub>=4.289**<br>**p<0.001**<br>(<0.001) | 2.41<br>(1.72;3.28) | 2.65<br>(1.94;3.49) | 0.331 |

Strong significant positive relationships were found in δ^15^N between blood cells and plasma (males: F_1,25_=18.846, P<0.001, r=0.656; females: F_1,33_=31.679, P<0.001, r=0.700; Figure 5), but not between feathers and blood cells (males: F_1,36_=0.036, P=0.850, r=0.032; females: F_1,45_=0.062, P=0.805, r=0.037). No significant positive relationship was found in residual δ^13^C between blood cells, plasma and feathers (all p’s>0.243).

**Figure 5.**
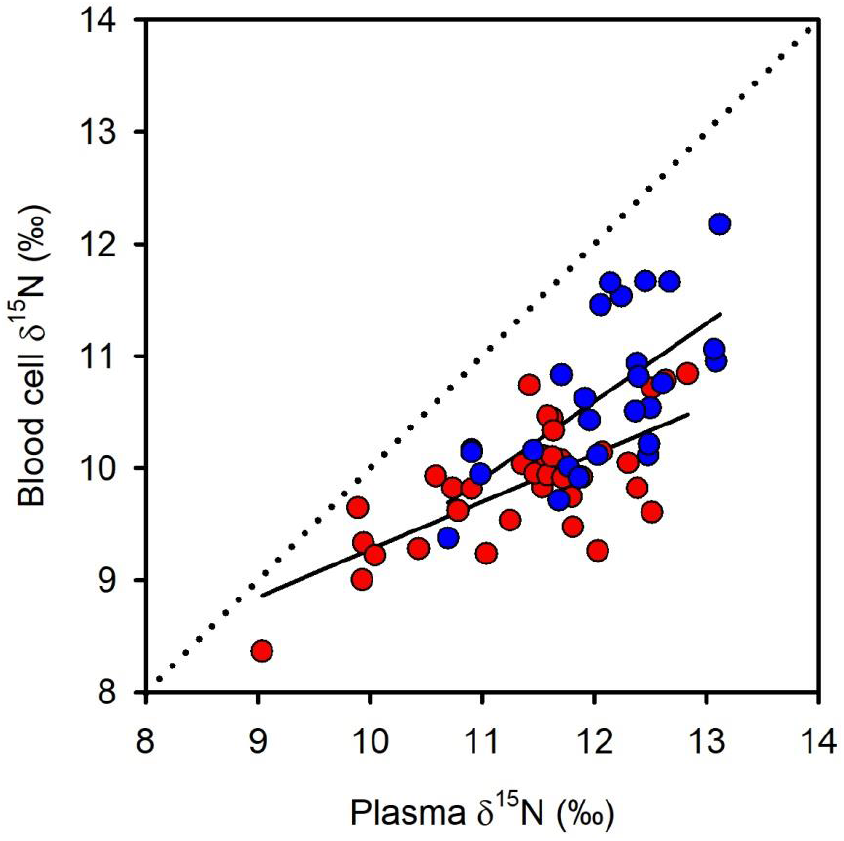
Relationships between blood cells and plasma δ^15^N values for male (n = 27, blue) and female (n = 35, red) snow petrels sampled at Ile des Pétrels, Adélie Land, East Antarctica. Males: F_1,22_=15.203, P<0.001, R^2^ = 0.409; females: F_1,20_=24.300, P<0.001, R^2^ = 0.549.

There was no significant relationship between isotopic values and body measurements or body condition (all p’s>0.08).

### Foraging trip performance and foraging efficiency

Foraging trip duration, length, speed and directions (Table 7), as well as mass gain and proportion mass gain (Table 8) did not differ between males and females. Foraging efficiency, measured as the proportionate daily mass gain while foraging, was significantly greater for females than for males (Table 8), and was greater for larger females with deeper bills (PC1: F_1,20_=5.279, P=0.033, r=0.457; bill depth: F_1,20_=8.630, P=0.008, r=0.549). In females, but not in males, foraging efficiency decreased with the duration of the foraging trip (Figure 6). Females were more efficient than males during short (<2 days) foraging trips, but for trips longer than 2 days, foraging efficiency was similar in males and in females (daily mass gain: t_39_=0.397, P = 0.693; proportion daily mass gain: t_39_=0.862, P = 0.394).

**Figure 6.**
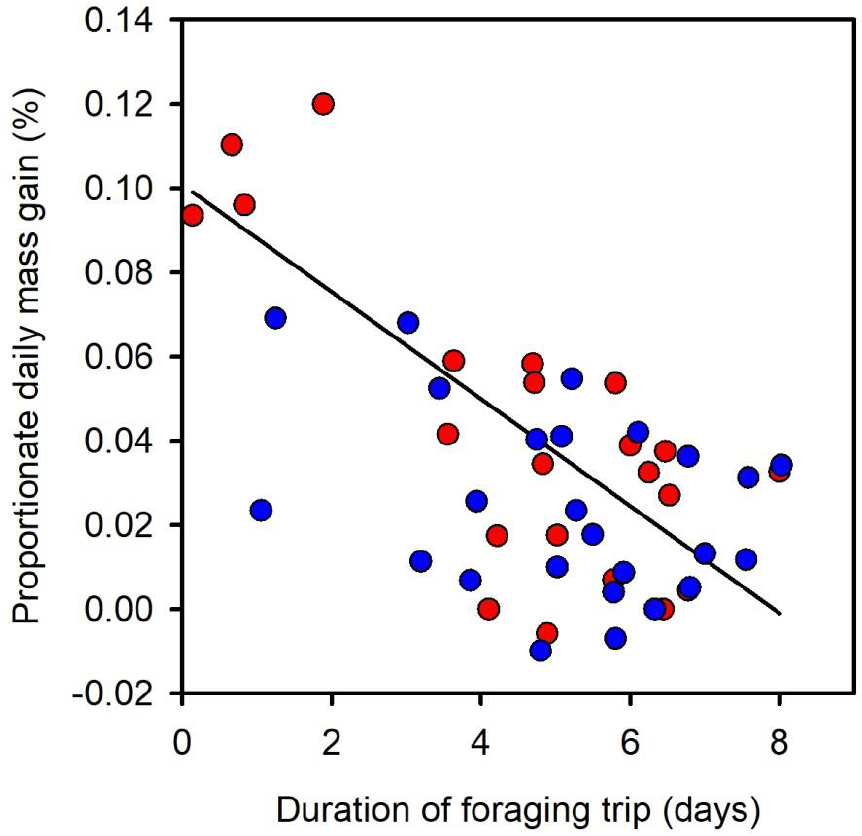
Foraging efficiency (proportionate daily mass gain while foraging) as a function of the total duration of the foraging trip for male (blue) and female (red and solid line) snow petrels sampled at Ile des Pétrels, Adélie Land, East Antarctica. For females: F_1,20_=25.349, P<0.001, R^2^ = 0.559.

**Table 7.** Summary of foraging trip metrics for snow petrels from Ile des Pétrels, Adélie Land, East Antarctica. For t-tests homogeneity of variances we checked using a Brown and Forsythe tests (Brown and Forsythe 1974), corrected p values are reported (uncorrected in brackets). Significant differences with a false detection rate of 0.10 are shown in bold.

|  | Sex | Mean (SD) | Range | t-test |
| --- | --- | --- | --- | --- |
| Trip duration (d) | Male | 5.2 (1.8) | 1.1-8.0 | $t_{45}=0.979$ |
| | Female | 4.8 (1.9) | 0.7-8.0 | $p=1.160$ (0.333) |
| Trip length (km) | Male | 851.6 (352.1) | 302.0-1957.5 | $t_{45}=0.292$ |
| | Female | 856.4 (389.0) | 173.8-1803.8 | $p=0.772$ (0.772) |
| Trip mean speed ($\text{km}\cdot\text{h}^{-1}$) | Male | 2.0 (0.6) | 1.2-3.5 | $t_{45}=0.762$ |
| | Female | 2.1 (0.4) | 1.7-3.2 | $p=0.788$ (0.450) |
| Trip maximum speed ($\text{km}\cdot\text{h}^{-1}$) | Male | 13.5 (2.6) | 9.2-21.4 | $t_{45}=0.670$ |
| | Female | 13.3 (2.5) | 9.5-18.9 | $p=0.709$ (0.506) |
| Trip start direction (°) | Male | 161.5 (114.0) | 0.8-355.2 | $t_{45}=1.066$ |
| | Female | 204.6 (143.2) | 2.6-348.6 | $p=2.045$ (0.292) |
| Trip mean direction (°) | Male | 190.9 (110.4) | 70.1-307.9 | $t_{45}=0.336$ |
| | Female | 209.2 (117.6) | 68.8-342.7 | $p=0.862$ (0.739) |
| Trip end direction (°) | Male | 167.5 (114.0) | 46.9-282.8 | $t_{45}=0.773$ |
| | Female | 184.8 (68.6) | 102.9-291.6 | $p=1.035$ (0.444) |

**Table 8.** Summary of metrics of foraging trip efficiency for snow petrels from Ile des Pétrels, Adélie Land, East Antarctica. For t-tests homogeneity of variances we checked using a Brown and Forsythe tests (Brown and Forsythe 1974), corrected p values are reported (uncorrected in brackets). Significant differences with a false detection rate of 0.10 are shown in bold.

|  | Sex | Mean (SD) | Range | t-test |
| --- | --- | --- | --- | --- |
| Mass at departure (g) | Male | 389.0 (30.4) | 331.5-464.0 | $t_{45}=6.014$ |
| | Female | 340.5 (24.0) | 300.0-385.0 | $p<0.001$ (<0.001) |
| Mass at return (g) | Male | 431.4 (34.0) | 365.0-495.0 | $t_{45}=4.467$ |
| | Female | 387.0 (33.9) | 320.0-460.0 | $p<0.001$ (<0.001) |
| BCI<sup>1</sup> at departure | Male | 377.1 (25.2) | 336.2-437.7 | $t_{45}=2.633$ |
| | Female | 358.1 (24.0) | 321.3-414.1 | $p=0.031$ (0.012) |
| BCI at return | Male | 419.5 (31.2) | 372.5-479.3 | $t_{45}=1.636$ |
| | Female | 405.0 (29.3) | 361.0-458.4 | $p=0.145$ (0.109) |
| $\Delta\text{mass}$ (g) | Male | 42.4 (38.1) | -20-110 | $t_{45}=0.392$ |
| | Female | 46.6 (34.6) | -10-95 | $p=0.697$ (0.697) |
| Daily mass gain ($\text{g}\cdot\text{day}^{-1}$) | Male | 9.2 (8.2) | -4.2-28.0 | $t_{45}=1.707$ |
| | Female | 14.6 (13.2) | -2.0-45.0 | $p=0.152$ (0.095) |
| Proportion mass gain (%) | Male | 0.11 (0.10) | -0.05-0.29 | $t_{45}=0.886$ |
| | Female | 0.14 (0.11) | -0.03-0.31 | $p=0.435$ (0.381) |
| Proportion daily mass gain (%) | Male | 0.02 (0.02) | -0.01-0.07 | $t_{45}=2.064$ |
| | Female | 0.04 (0.04) | -0.01-0.12 | $p=0.089$ (0.045) |

<sup>1</sup> Body Condition Index

### Regulation of the foraging trips

To investigate how birds regulate foraging trips according to the depletion of their body reserves, we correlated the body condition at departure with the duration of the foraging trips and the mass gain metrics while foraging. Foraging trip duration was not correlated to body condition at departure (Pearson correlation coefficient: p = 0.417 for females, p = 0.576 for males), but mass gain and proportionate mass gain were negatively related to body condition at departure for both sexes (all P’s<0.005; Figure 6). In addition, in males, but not in females, daily mass gain and proportionate daily mass gain were negatively correlated to body condition at departure (males: all P’s<0.010; females: all P’s>0.429; Figure 7). Male (but not female) body condition at return from a foraging trip was positively correlated to the time spent at sea (Pearson correlation coefficient: p = 0.05).

**Figure 7.**
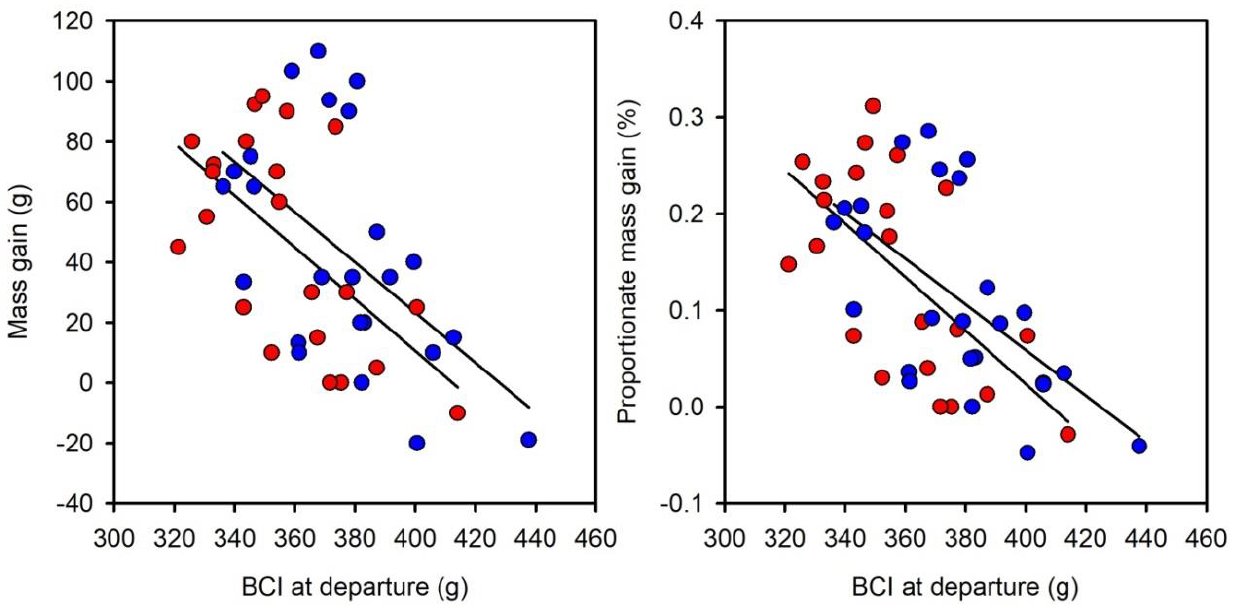
Mass gain and proportionate mass gain as a function of body condition before a foraging trip for male (blue) and female (red) snow petrels sampled at Ile des Pétrels, Adélie Land, East Antarctica. Male mass gain: F_1,23_=10.010, P=0.004, r^2^=0.303; female mass gain: F_1,20_=11.071, P=0.003, r^2^=0.356; male proportionate mass gain: F_1,23_=12.361, P=0.002, r^2^=0.350; female proportionate mass gain: F_1,20_=13.258, P=0.002, r^2^=0.399.

### Factors affecting mass gain at sea

For females, but not for males, mass gain, proportion mass gain and body condition at return from a foraging trip were positively correlated to mean and maximum sea ice concentration of the foraging trip locations (females: all P<0.049; males: all P>0.232; Figure 8). For males and females, there was no relationship between bathymetry and mass gain, proportion mass gain, and body condition at return (all P>0.100).

**Figure 8.**
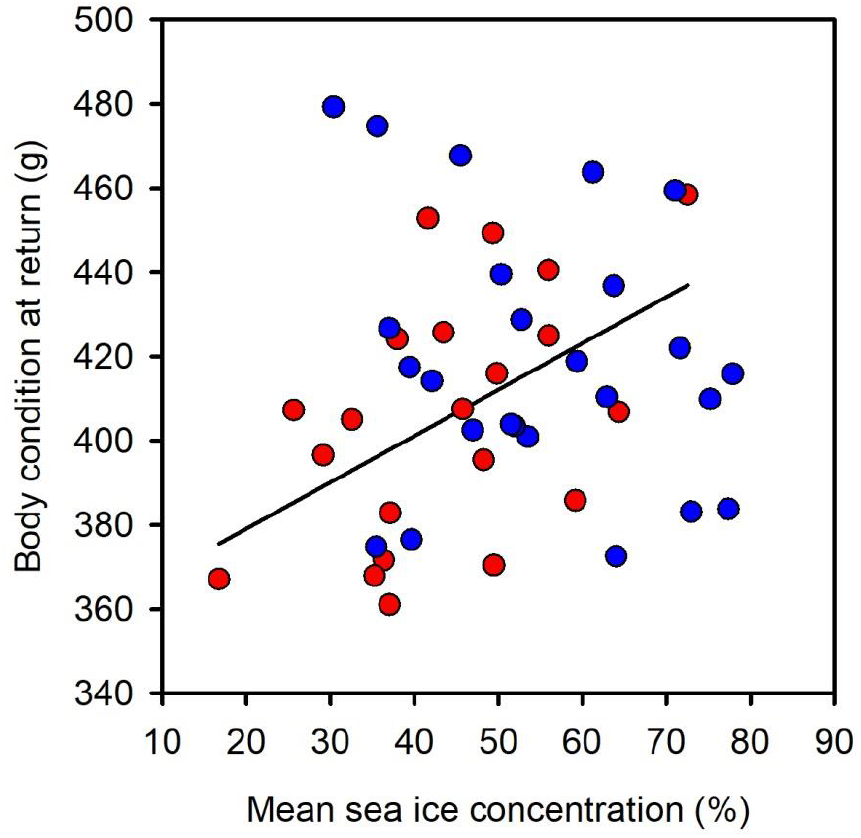
Body condition at return from a foraging trip as a function of the mean sea ice concentration of the foraging trip locations for male (blue) and female (red and solid line) snow petrels sampled at Ile des Pétrels, Adélie Land, East Antarctica. For females: F_1,19_=6.106, P=0.023, R^2^ = 0.243.

## Discussion

This study provides clear evidence of sexual segregation and foraging tactics in snow petrels. In accordance with our prediction, we found evidence for sexual segregation in diet, with males feeding on average on higher trophic level prey when compared to females, but no evidence for spatial segregation as indicated by spatial data and δ^13^C isotopic data. Males and females differed in their usage of sea ice, providing evidence for sex-specific habitat segregation.

### Differences in habitat use

During incubation males and females foraged predominantly in pack-ice areas over the deep Antarctic continental shelf and adjacent continental margin (500-900 m, due to the isostatic effect of the ice sheet), and to a lesser extent in oceanic waters. These results are consistent with previous observational work at sea showing that high densities of breeding snow petrels in the Ross Sea were found within the pack ice along the continental slope (Ainley et al. 1984). The low tissue δ^13^C values of snow petrels is a consistent characteristic of consumers foraging in high-Antarctic waters (Cherel 2008, Cherel et al. 2011). Blood cell, plasma and feather δ^13^C values were similar in males and females, which indicates that both sexes foraged offshore in pelagic waters without an obvious neritic-oceanic δ^13^C gradient in high-Antarctica (Cherel et al. 2011). Blood cell and plasma δ^13^C values of birds from Adélie Land were similar to values obtained from snow petrel muscle tissue in the Weddell Sea (Rau et al. 1992) between 64°S and 66°S, but were slightly lower than those measured in whole blood of birds from Hop Island (Hodum & Hobson 2000). However, the shape of the relationships between foraging intensity and SIC suggested that males and females used different sea ice habitats. Female foraging intensity was highest for SIC between ≈20% and ≈40%, and then decreased non-linearly for higher SIC, with a sharp decrease for SIC higher than ≈70%. By contrast male foraging intensity remained high for high SIC. Therefore, although foraging intensity decreased with increasing SIC for both sexes, males foraged more intensively in high sea ice concentration areas (> 70%) than females. Males and females made greater use of pack-ice areas over the continental shelf and continental margin than of oceanic pack-ice areas, but males were more likely to forage and foraged more intensively on the continental margin (−550 to −950 m) than females.

Few studies have simultaneously quantified between-sex differences in habitat use and foraging behavior in marine species in relation to dynamic oceanographic features such as sea ice. In the northern gannet (*Morus bassanus*), sexual segregation was driven largely by spatial and habitat segregation with males, smaller than females, mainly foraging in coastal mixed waters where net primary production was high, and females mainly foraging in offshore stratified waters (Cleasby et al. 2015). Similarly, the sex-specific habitat use reported in the monomorphic Barau’s petrel (*Pterodroma baraui*) during the prelaying exodus (males used more frequently marine areas with high productivity) can be partly explained by spatial segregation between sexes (Pinet et al. 2012). During the incubation and chick rearing period, they did not find evidence for habitat segregation and foraging areas largely overlapped. In the Adélie penguin (*Pygoscelis adeliae*) at Pointe Géologie, females foraged more intensively in areas of higher sea ice concentration than males during the guard stage, and there was spatial segregation between sexes with females foraging further from the colony than males (Widmann et al. 2015). Using a multiyear comprehensive dataset, Paiva et al. (2017) found that sexual segregation in foraging areas and foraging habitats of Cory’s shearwaters (*Calonectris borealis*) varied between years, with greater sexual (habitat and spatial) segregation during years when sea surface temperatures were higher and chlorophyll *a* concentrations were lower, presumably corresponding to lower food availability. In favorable years no spatial segregation was observed and habitat segregation was low. The hatching success of snow petrels during the 2015/2016 breeding season was 46.9%, i.e. lower than the longterm average of 63.3% (Chastel et al. 1993), suggesting that environmental conditions were relatively poor. However, we did not observe spatial segregation between the sexes but foraging habitat use differed, with males foraging more frequently in high sea ice concentration areas than females. Such a pattern was found in the wandering albatross (*Diomedea exulans*) at South Georgia in which, despite no clear sexual segregation at large scales, sex-specific microhabitat selection was found during the chick-rearing period, resulting in sexual segregation in core foraging areas (Pereira et al. 2018). Multiple years of tracking are needed to shed light into the effects of environmental stochasticity (sea ice variability) on habitat segregation and spatial segregation.

As opposed to other highly sexually size-dimorphic seabirds (wandering albatross: Weimerskirch et al. 1993, giant petrels *Macronectes spp*.: Gonzáles-Solís et al. 2000, boobies *Sula spp*.: Weimerskirch et al. 2009, frigatebirds *Fregata spp*.: Hennicke et al. 2015) snow petrels did not show spatial segregation in their foraging habitat during incubation. Spatial segregation in snow petrels may occur during other periods of the year such as during the chick-rearing period when which food requirements are particularly high for provisioning the chick. Alternatively, this lack of spatial segregation may be constrained by the specific foraging habitat requirements of snow petrels. These seabirds forage exclusively in a sea ice environment, which is limited during the breeding season around breeding colonies and may thus constraint males and females to spatially overlap at a broad spatial scale.

### Influence of sex on diet and foraging tactics

The snow petrel diet is relatively well known during the chick-rearing period and isotopic data together with prey biometric data suggest that snow petrels mainly feed on postlarvae and juvenile Antarctic silverfish (*Pleuragramma antarcticum*) (Ridoux & Offredo 1989, Hodum & Hobson 2000, Pinkerton et al. 2013). Although, snow petrel diet during incubation remains poorly known, δ^15^N values obtained in our study are similar or slightly higher than those found in other studies during the chick rearing period (Hodum & Hobson 2000, Delord et al. 2016), suggesting a similar diet. Nevertheless, and despite large overlap in their core isotopic niches as indicated by the standard ellipse areas, female snow petrels had lower δ^15^N values than males for all tissues sampled, which suggests they were feeding on lower trophic level prey than males. Similar results were found by Tartu et al. (2014) for blood cells during the pre-laying period. We speculate that there might be at least two reasons for this. First, compared to males, females may feed more frequently on other prey than Antarctic silverfish, such as crustaceans which are situated at a lower trophic level than Antarctic silverfish. Indeed, diet studies indicate that snow petrels also feed on crustaceans such as *Euphausia superba, E. crystallorophias, Themisto gaudichaudii*, and other amphipods (Ainley et al. 1984, Ridoux & Offredo 1989) which have lower δ^15^N values than Antarctic silverfish (Pinkerton et al. 2013). Second, females may feed on Antarctic silverfish in similar proportions than males but on smaller sized individuals (i.e. younger). It is known that δ^15^N values increase with body length (and age) in Antarctic silverfish from ≈7-8‰ in larvae (10-20 mm standard length) to ≈10-11‰ in juvenile and adult fish (Giraldo et al. 2011, Pinkerton et al. 2013). It is currently unknown whether sea ice concentration and characteristics differentially affect the spatial distribution of Antarctic silverfish age-classes. However, it is likely that females fed more on crustaceans than on young silverfish since crustaceans have much lower δ^15^N values than young silverfish (Cherel 2008). Thus, our results suggest that males ate more silverfish in areas with higher sea ice concentration.

The strong positive relationship between plasma δ^15^N and blood δ^15^N indicates short term (over weeks) consistency in trophic level between successive foraging trips during incubation. Values of δ^15^N in plasma and feathers did not differ in both sexes (Appendix 1), but blood δ^15^N were smaller than feather and plasma δ^15^N in both sexes, suggesting that males and females fed on lower trophic level prey prior to incubation than during the breeding season. Short and long term consistency in foraging water masses was also low as indicated by the lack of relationship between plasma and blood δ^13^C, and between feather and blood δ^13^C, respectively. Indeed tracking data indicated that birds foraged on the continental shelf, continental margin, and to a lesser extent in oceanic waters. Values of δ^13^C in feathers were higher than those in blood and plasma for both sexes (Appendix 1), suggesting that during the latter part of the breeding season and the beginning of the non-breeding season snow petrels foraged in more oceanic waters (snow petrels start molting during the chick rearing period and until early May (Beck 1970, Delord et al. 2016). This period coincides with the sea ice growth and its northward extension.

The negative relationship between mass gain (and proportion daily mass gain) during a foraging trip and body condition at departure for a foraging trip (i.e. at the end of fasting while incubating the egg), indicated that males and females were able to regulate their body reserves as found in other Procellariiform species (Chaurand & Weimerskirch 1994, Gonzáles-Solís et al. 2000). Although both sexes regulated body condition, this ability seemed greater for females than for males. Indeed, body condition at departure for a foraging trip was lower in females than in males, but similar for both sexes at return from a foraging trip despite similar trip durations. This is further supported by the fact that females had higher daily mass gains and proportion daily mass gains than males. However, this greater ability in females may be partly explained by the fact that females undertook short foraging trips during which mass gain was particularly high (Figure 1). Although some males also made short foraging trips, mass gain was still lower than female mass gain during these trips. Therefore, these results suggest that female foraging efficiency was similar in males and females, except during short (<2 days) foraging trips during which females appeared more efficient. We suspect that some females undertook short foraging trips during their incubation shift in order to restore their body condition to avoid abandoning the egg while their partner was foraging at sea. This could result from the lower fasting capacities of females compared to males due to their smaller body size (Barbraud & Chastel 1999).

Interestingly, the ability of females (but not of males) to restore their body condition during a foraging trip was affected by sea ice concentration. Indeed, female body condition at return from a foraging trip was positively related to sea ice concentration in the foraging area, contrary to males. This suggests that areas with heavy sea ice concentration were more profitable. This is further supported by the positive relationship between male (but not female) body condition at return from a foraging trip and time spent at sea, and given that males foraged more frequently in high sea ice concentration areas. Thus, foraging on highly nutritional preys such as silverfish in high sea ice concentration areas might be more efficient to restore body condition that feeding in more open water areas.

Body condition at the start of a foraging trip was not related to the time spent at sea, suggesting that the time spent at sea was not only dependent on the restoration of body condition. Although only a few birds returned to undertake the next incubation shift after losing mass (n = 3, 6.3%) or without gaining mass (n = 3, 6.3%), this suggests that mass gain alone does not explain the decision to return to the colony. Perhaps birds took into account the increased probability of partners deserting the egg with the increasing duration of the foraging trip (Tveraa et al. 1997).

Thus, incubating female snow petrels seemed more efficient at restoring their body condition during a foraging trip despite similar trip duration, length or speed, while foraging areas were identical to those of males at a broad spatial scale. However, this higher efficiency mainly concerned short (<2 days) foraging trips. In addition, our results show that females foraging in high sea ice concentration areas foraged more efficiently (this relationship holds when excluding foraging trip <2 days), and female fed on lower trophic level preys that males. Together, these results suggest that areas with high sea ice concentration may be more profitable for resource acquisition, perhaps due to higher abundance, availability or quality of prey such as the Antarctic silverfish.

### Factors underlying sexual segregation

Sex differences in foraging behavior could result from the influence of sexual size dimorphism on foraging efficiency and intra-specific competition (forage-selection hypothesis and scrambled competition hypothesis). The positive relationship between female bill depth and proportion daily mass gain suggests that foraging efficiency is size dependent in females, which are smaller than males. Our results also suggest that the most favorable areas were areas of high sea ice concentration (females body condition at return increased with increase sea ice concentration, male body condition at return increased with foraging trip length), which were used less frequently by females. Therefore, it is possible that females were excluded from high sea ice concentration areas via direct competition. This could possibly indicate that male and female snow petrels try to avoid competition and thus diverged in habitat preference in more profitable areas, where intra-specific competition might be more intense. Such a mechanism was also proposed to explain sex-specific differences in broad scale foraging areas in highly sexually size dimorphic species (wandering albatross: Weimerskirch et al. 1993, Shaffer et al. 2001; giant petrels: Gonzáles-Solís et al. 2000), but also in foraging habitat at a microhabitat scale (Pereira et al. 2018). A major assumption of the intersexual competition hypothesis is that prey capture should be a function of bill size (Selander 1966, Shine 1989). Although we do not have the data in hand to test this prediction explicitly, we note that δ^15^N values suggested that females consumed lower sized prey than males (crustaceans vs fish). Females with thicker bills were also more efficient during their foraging trip, suggesting they were feeding on more profitable prey, and bill size was among the most sexually dimorphic phenotypic trait in this species.

Sex-specific niche divergence and habitat segregation can also arise from a difference between sexes in parental roles and investment (the activity budget hypothesis, Clarke et al. 1998, Thaxter et al. 2009, Weimerskirch et al. 2009, Pinet et al. 2012). Although males undertake a greater investment in chick provisioning though higher feeding frequencies (Barbraud et al. 1999), there is little differentiation in the reproductive role of male and female snow petrels during incubation. Males make slightly shorter foraging trips than females during incubation (Isenmann 1970, Barbraud et al. 1999), but in average the total time spent foraging during the incubation period is very similar for both sexes (males: average 19.8 days, females: average 21.0 days, Barbraud 1999), indicating that the roles of male and female snow petrels do not appear to differ substantially during incubation. Therefore, it seems unlikely that such limited constraints related to reproductive role specialization could explain why female snow petrels foraged less intensively in high sea ice concentration areas; this hypothesis can probably, therefore, be discounted. Sex-specificity in flight performance may also be responsible for sexual segregation (Shaffer et al. 2001, Phillips et al. 2004). Indeed, sexual dimorphism in wing area and wing loading in several albatross species may partially explain large-scale sexual segregation in foraging areas in these species: sex-specific foraging locations were likely influenced by activity budgets since smaller birds are more efficient flyers. Therefore, other aspects of the morphology not measured here, such as wing loading and agility, may be important. Female snow petrels appear to have a lower aspect ratio and lower wing loading than males (Spear & Ainley 1998), suggesting they might be less flight efficient but more maneuverable than males. However, since there was no spatial segregation between sexes at large-spatial scales, environmental conditions potentially affecting flight efficiency (wind speed) were identical for males and females. Thus, these aspects are also unlikely to be of importance in snow petrels to explain sex-specific foraging habitat use during incubation.

Overall, our study demonstrates sex-specific foraging tactics in a highly sexually size dimorphic species during the incubation period, probably driven by intra-specific competition. Results indicate an absence of sexual segregation at a broad-spatial scale, but suggest that sexual segregation in snow petrels is mediated by habitat segregation at a microhabitat scale. Males foraged more intensively than females in high sea ice concentration areas, which seemed to be more profitable in terms of resource acquisition as results suggest that males ate more fish in these areas. Studying sex-specific foraging tactics during the entire breeding period, thus including the pre-laying exodus and the chick-rearing period, is however necessary to better understand the underlying drivers of sexual segregation in snow petrels and in marine predators in general (Pinet et al. 2012). Sexual segregation in foraging behavior may also vary between years as a function of environmental conditions (Cleasby et al. 2015, Paiva et al. 2017), highlighting the need for multi-year tracking studies.

## Data accessibility

Tracking data are available on MoveBank https://www.movebank.org/.

## Acknowledgements

This preprint has been peer-reviewed and recommended by Peer Community In Ecology (https://dx.doi.org/10.24072/pci.ecology.100025). The authors thank G. Brodin for his help in designing GPS loggers and extracting some GPS data, J. Vasseur who helped with deploying and recovering some GPS loggers and collecting tissue samples. The authors are grateful to the analytical Plateformes of the LIENSs laboratory for the access to their analytical facilities, and in particular to M. Brault-Favrou and G. Guillou for running stable isotope analyses. The work was supported financially and logistically by the French Polar Institute (IPEV program No. 109, PI H. Weimerskirch), the French Southern Territories administration (TAAF). This study is a contribution to program SENSEI funded by the BNP Paribas Foundation. The field study was approved by the IPEV ethics committee and Comité de l’Environnement Polaire. The IUF (Institut Universitaire de France) is acknowledged for its support to PB as a Senior Member. We thank Dries Bonte and an anonymous reviewer for helpful suggestions on the manuscript.

## Appendix

**Appendix I.** Testing for differences in δ^15^N (‰) and δ^13^C (‰) values between tissues for male and female snow petrels sampled Ile des Pétrels, Adélie Land, East Antarctica. t indicates Student’s t-tests with df, Z indicates Wilcoxon rank test. ** indicates P < 0.01, *** indicates P < 0.001 after applying the Benjamini-Hochberg procedure with a false discovery rate of 0.10. Values above diagonal are for δ^15^N (‰), values below diagonal are for δ^13^C(‰). δ^15^N (‰) and δ^13^C (‰) values in feathers were corrected following Cherel et al. (2014a) before comparison with blood cells. δ^13^C (‰) values for plasma were normalized following Post et al. (2007) and Cherel et al. (2014b).

|  | Plasma | Blood | Feather |
| --- | --- | --- | --- |
| **Male** |  |  |  |
| Plasma | - | $t_{47}=7.206^{***}$ | $t_{47}=0.008$ |
| Blood cells | $Z=2.743^{**}$ | - | $t_{47}=3.060^{**}$ |
| Feather | $t_{47}=17.166^{***}$ | $t_{47}=29.033^{***}$ | - |
| **Female** |  |  |  |
| Plasma | - | $t_{42}=0.039$ | $t_{42}=0.435$ |
| Blood cells | $t_{42}=0.039$ | - | $t_{42}=2.967^{**}$ |
| Feather | $t_{42}=23.219^{***}$ | $Z=4.107^{***}$ | - |

